# cytoKernel: Robust kernel embeddings for assessing differential expression of single cell data

**DOI:** 10.1101/2024.08.16.608287

**Authors:** Tusharkanti Ghosh, Ryan M Baxter, Souvik Seal, Victor G Lui, Pratyaydipta Rudra, Thao Vu, Elena WY Hsieh, Debashis Ghosh

## Abstract

High-throughput sequencing of single-cell data can be used to rigorously evlauate cell specification and enable intricate variations between groups or conditions. Many popular existing methods for differential expression target differences in aggregate measurements (mean, median, sum) and limit their approaches to detect only global differential changes. We present a robust method for differential expression of single-cell data using a kernel-based score test, cytoKernel. cytoKernel is specifically designed to assess the differential expression of single cell RNA sequencing and high-dimensional flow or mass cytometry data using the full probability distribution pattern. cytoKernel is based on kernel embeddings which employs the probability distributions of the single cell data, by calculating the pairwise divergence/distance between distributions of subjects. It can detect both patterns involving aggregate changes, as well as more elusive variations that are often overlooked due to the multimodal characteristics of single cell data. We performed extensive benchmarks across both simulated and real data sets from mass cytometry data and single-cell RNA sequencing. The cytoKernel procedure effectively controls the False Discovery Rate (FDR) and shows favourable performance compared to existing methods. The method is able to identify more differential patterns than existing approaches. We apply cytoKernel to assess gene expression and protein marker expression differences from cell subpopulations in various publicly available single-cell RNAseq and mass cytometry data sets. The methods described in this paper are implemented in the open-source R package cytoKernel, which is freely available from Bioconductor at http://bioconductor.org/packages/cytoKernel.

## 1 Introduction

Technological advancements have revolutionized the field of high-throughput single-cell sequencing (sc-seq) (Wen et al., 2022; Tiberi et al., 2022), particularly through single-cell RNA sequencing (scRNA-seq) (Saliba et al., 2014) and advanced high-dimensional (flow or mass) cytometry (Pyne et al., 2009). Single-cell sequencing (sc-seq) data have enabled an in-depth exploration of biological processes at the single-cell level (Stuart and Satija, 2019; Papalexi and Satija, 2018) The single-cell sequencing approach exceeds the capabilities of conventional bulk analysis by providing spatial-temporal insights into biological processes with high resolution (Chen, 2022). Sc-seq plays a pivotal role in elucidating cellular heterogeneity, detecting rare cell subpopulations, isolating targeted biomarkers, and profiling distinctive molecular characteristics at the single cell level (Ren et al., 2018; Giladi and Amit, 2018).

A widely adopted approach in sc-seq for unraveling both intrinsic and extrinsic biological processes involves identifying genes (for scRNA-seq) or proteins (for high-dimensional cytometry) that exhibit differential expression (DE) (Zhang and Guo, 2022). Differential expression (DE) method facilitate the isolation and detailed examination of specific signals emanating from a particular cell subpopulation of interest. However, challenges arise due to the high heterogeneity and prevalence of zero counts in sc-seq data, complicating the statistical modeling and analysis (Kharchenko, 2021; Luecken and Theis, 2019).

Classical methods for analyzing differential expression (DE), traditionally applied to bulk RNA-seq data, have also been adapted for single-cell measurements, as indicated in studies by Soneson and Robinson (2018); Wang et al. (2019); Chen et al. (2020); Squair et al. (2021). These methods have been shown to be effective on aggregated data by summarizing single-cell sequences into aggregate counts, also referred to as pseudo-bulk (PB) approach (Crowell et al., 2020), thus allowing the application of bulk RNAseq analysis methods. Traditional DE methods, particularly those based on pseudo-bulk data analysis, often focus on aggregating single-cell data to represent a ‘bulk’ sample for each subject, thereby potentially overlooking intricate inter-subject variations in gene expression. In response to these complexities, several methods have been specifically developed for DE analysis in the context of single-cell data (Tiberi et al., 2022; You et al., 2023). These include scDD (Korthauer et al., 2016), SCDE (Kharchenko et al., 2014), BASiCS (Vallejos et al., 2015; Eling et al., 2018), and mixed models (Tung et al., 2017; Wills et al., 2013). Mixed-effects models, like the hurdle model described by Finak et al. (2015) and analyzed by Velmeshev et al. (2019) in autism research, are also commonly used. The hurdle model in MAST can be fit with a mixed-effects model (Finak et al., 2015). The model incorporates fixed effects for variables like case/control status and random effects at the subject level which specifies a logistic regression model for the expression rate and a linear model for the logarithmic non-zero expression. Despite their innovations, these methods exhibit certain limitations. For example, BASiCS does not cater to cell-type specific differential testing between conditions, scDD struggles with covariates and biological replicates, and others like PB, SCDE, MAST, and mixed models have shown limited efficacy in detecting differential patterns beyond mean differences in previous studies (Korthauer et al., 2016; Crowell et al., 2020). Conducting reliable statistical inference with MAST, such as testing for differential expression (DE) at a specified level of significance, within the framework of a fitted mixed-effect model, presents notable challenges. As detailed in (Zhang and Guo, 2022), the incorporation of random effects often renders conventional likelihood ratio tests unsuitable in these contexts. Furthermore, the practical application of the hurdle model is constrained by its parametric assumptions, which might not be representative of real single-cell datasets.

To bridge this analytical gap, we introduce cytoKernel, a methodology for generating robust kernel embeddings via a Hilbert Space approach designed for sc-seq data analysis. CytoKernel diverges from traditional methods by conceptualizing the cell type-specific gene expression of each subject as a probability distribution, rather than as a mere aggregation of single-cell data into pseudo-bulk measures. This probabilistic approach allows for a more sophisticated and nuanced comparison of gene expression across subjects, capturing complexities that pseudo-bulk methods may miss, such as subtle shifts and patterns in expression not observable through average measures alone.

## 2 Materials and Methods

### 2.1 Overview of the semiparametric logistic regression model via Functional Hilbert space

#### 2.1.1 cytoKernel-PB: A Pseudo Bulk Approach

Traditional pseudo bulk-based differential expression analysis usually involves the assessment of individual features within cell subpopulations through a parametric model (Love et al., 2014; Robinson et al., 2010). This approach includes analyzing each feature by aggregating cells from each subject which generates a p-value. Then, a feature is declared significant, if its p-value is lower than a particular threshold, which is often corrected for multiple testing (Benjamini and Hochberg, 1995; Benjamini and Yekutieli, 2001).

For a fixed number of cells within a subject *i*, we observe the data triplet {**z**_*i*_, **x**_*i*_, *y*_*i*_}, *i* = 1, …, *n* where *n* is the total number of subjects obtained from a case control study for gene *g* (omitting the index *g* = 1, …, *G* for simplicity). For subject *i, y*_*i*_ is the case control label taking values either 0 (control) or 1 (case), **x**_*i*_ is a *q* × 1 vector of covariates, **z**_*i*_ is a *c*_*i*_ × 1 single cell vector. *c*_*i*_ denotes the number of cells at single cell resolution for subject *i*. In other words, the dimension of **z**_*i*_ equals the size of the single cell vector for subject *i*.

A key element for the pseudo bulk approach is the reduction of the single cell vector **z**_*i*_, which varies in dimension according to the single cell count *c*_*i*_ for each subject *i*, into a scalar quantity,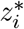. This reduction is a critical step in aggregating single cell data into a pseudo-bulk format for the data. We can achieve this using various aggregation techniques such as averaging, median calculation, or pooling of data (Crowell et al., 2020). These aggregation methods are carefully selected to balance the simplification of data with the preservation of crucial biological signals (Finak et al., 2015).

Continuing with the model development for the pseudo bulk approach, we assume that an intercept is included in **x**_*i*_. The binary outcome *y*_*i*_ depends on **x**_*i*_ and 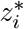 through the following semiparametric logistic regression model:

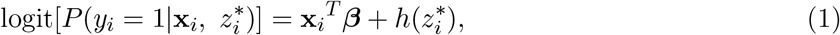

where ***β*** is a *q* × 1 vector of regression coefficients, and 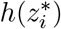 is an unknown centered smooth function.

Model (1) is semiparametric in the sense that it does not put any assumptions on *h*(·) except that it is assumed to lie in a certain functional space ℋ_*k*_. The covariate effects are modeled parametrically, while the pseudo bulk scalar quantity 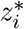 is modeled non parametrically. A non parametric assumption for *h*(·) reflects our limited knowledge the functional forms of the specific gene *g*. Note that, when 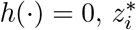 has no association with the group labels *y*_*i*_. Hence, a differentially expressed feature will lead to a rejection of the null hypothesis *h*(·) = 0. Note, if 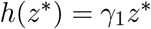,for any arbitrary *γ*_1_, model (1) becomes the generalized linear model (Goeman et al., 2011).

#### 2.1.2 cytoKernel-sc: A Comprehensive Single-Cell Approach Capturing Full Distributional Characteristics

For the sake of simplicity, we use the same notation from the previous section used in describing the data triplets {**z**_*i*_, **x**_*i*_, *y*_*i*_}, *i* = 1, …, *n* where *n* is the total number of subjects obtained from a case control study for gene *g*. The following model is considered for the analysis of differential expression of each feature in the cytoKernel methodology:

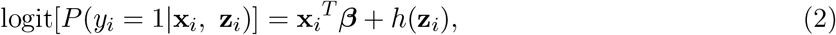

where *h*(·) is a centered smooth function in a RKHS (reproducing kernel Hilbert space; Wahba (1990)) spanned by kernel *k* and **z**_*i*_ is the single cell vector with dimension *c*_*i*_ × 1. Note, (2) is assumed to lie in ℋ_*k*_, the Hilbert space. Such kernel-based models are more robust to issues lies like model misspecification. Similar to the hypothesis in the cytoKernel-PB section, a differential expressed gene will lead to the rejection of the null hypothesis *H*_0_ : *h*(·) = 0.

### 2.2 Kernel-based Score test

We provide a detailed implementation of the Kernel-based score test for sc-seq datasets, particularly focusing on the cytoKernel-PB version. The key adaptation for the cytoKernel-sc version involves substituting the variable 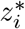 of the cytoKernel-PB version with **z**_*i*_ in the the Kernel-based score test framework. Apart from this specific modification, all other steps in the procedure remain consistent with the kernel-based semiparametric logistic regression model for cytoKernel-sc. This approach upholds the methodological strictness of the Kernel-based score test, while incorporating the specific modifications using the distance metric required for its effective application to single-cell datasets.

The RKHS ℋ_*k*_, is generated by a positive definite kernel function *k*. The statistical properties of ℋ_*k*_ imply that any function *h*(*z* can be written as a linear combination of given function *k*(·, ·). Let *K* bet the *n*×*n* Gram matrix with 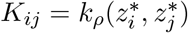 is the reproducing kernel of the RKHS which contains *h*(·), and *ρ* is an unknown kernel parameter.

A link between kernel machine regression and linear mixed models was established in the context of the semiparametric modeling for high dimensional data (Liu et al., 2008).

Assuming that *h*(·) lies within a RKHS, *h*(·) ∈ ℋ_*k*_, ***β*** and *h*(·) can be simultaneously estimated by maximizing the penalized log-likelihood function

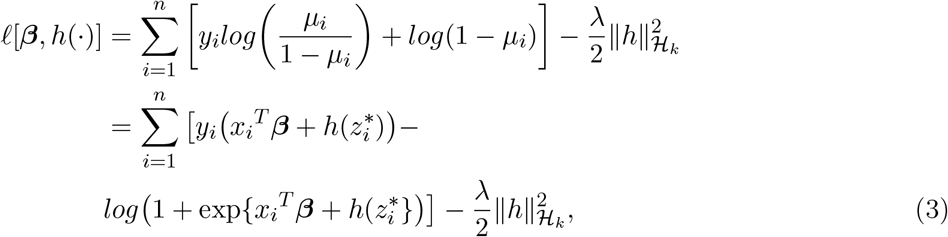

where 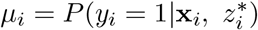 and *λ* is a regularization parameter that contributes to the the balance between model complexity and goodness of fit (Liu et al., 2008). If *λ* = 0, it reflects a saturated model at its boundaries whereas *λ* = 0 reduces the model to a fully parametric logistic regression model. There are two unknown parameters in *ℓ*[***β***, *h*(·)], the regularization parameter *λ* and bandwidth parameter *ρ*. We control the magnitude of the unknown function *h*(·) using *λ*. Meanwhile, *ρ* controls the smoothness of *h*(·) (Liu et al., 2008). The choice of *ρ* has a strong influence on the resulting estimate, so choosing an optimal value of *ρ* is critical.

According to Liu et al. (2008), it is possible to approach *ℓ*[*β, k*(·)] from a generalized linear mixed models (GLMM) perspective. As logistic regression is a special case of GLMM, the kernel estimator within the semiparametric logistic regression model parallels the penalized quasi-likelihood function from a logistic mixed model, letting *τ* = 1*/λ* denote the regularization parameter and *ρ* the bandwidth parameter (Liu et al., 2008; Zhan et al., 2015; Jensen et al., 2019). These parameters can be treated as variance components, where *k*(·) ∼ *N* (0, *τ K*(*ρ*)) can be treated as a subject-specific random effect and the covariance matrix *K*(*ρ*) is an *n* × *n* kernel matrix (Liu et al., 2008; Carpenter et al., 2021). This means that estimating *β* and *k*(·) can be done by maximizing the penalized log likelihood:

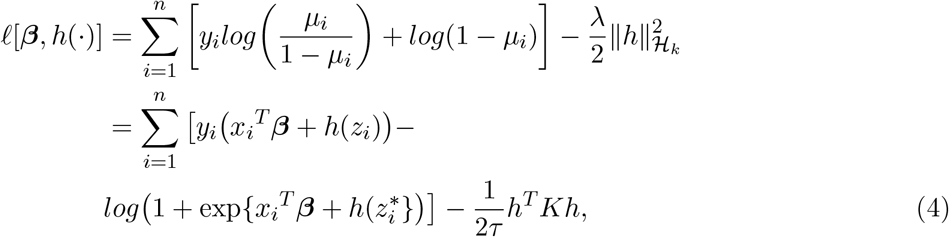

where *h* = *Kα* and *τ* = 1*/λ* (Liu et al., 2008). This provides an advantage as it allows for testing of the null hypothesis *H*_0_ : *τ* = 1*/λ* = 0 without explicit specification of basis functions. The function *h*(·) can then be understood as subject-specific random effects with mean 0 and variance *τ K*_*ρ*_. Testing for an association between binary outcome and the distribution of features is then equivalent to testing the null hypothesis

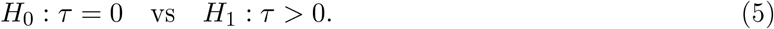

We use the modified kernel association test from Chen et al. (2016), tailored for small sample sizes, which is frequently applied in various gene expression and microbiome analyses including metabolomics studies. The standard quadratic score statistic for kernel association tests is given by:

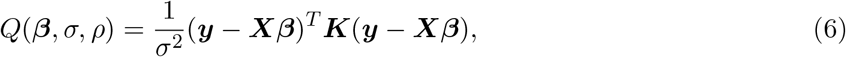

where ***y*** = (*y*_1_, *y*_2_, …, *y*_*n*_)^*T*^ and ***X*** = (***x***_1_, ***x***_2_, …, ***x***_*n*_)^*T*^. To account for high variability in the estimates of *σ*^2^ with smaller sample sizes, adjustments are made. The null distribution of *Q* is then approximated as a weighted sum of *χ*^2^ distributions using the Davies method (Davies, 1980).

#### 2.2.1 Gaussian Kernel and empirical choice of the tuning bandwidth parameter

Let 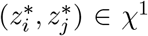 be two be two arbitrary Pseudo-Bulk (PB) gene expression measurements, where *χ*^1^ ≡ ℝ. We define a distance based Gaussian Kernel based on cytoKernel-PB,

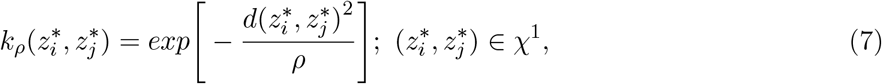

where 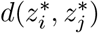 denotes the distance metric (square root of the Euclidean (*L*_2_) norm between pairwise Pseudo-Bulk (PB) gene expression measurements for subjects, *i* and *j*, respectively, and *ρ >* 0. The Gaussian kernel is employed, with the median of pairwise Euclidean distances between all 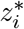 and 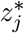 serving as an empirical estimate for the bandwidth parameter *ρ*. The selection of the Gaussian kernel is driven by its characteristic property, ensuring that the embedding of probability measures through the kernel function yields unique representations. Equation (7) is a well-defined kernel is shown in Zhan et al. (2015).

### 2.3 Employing divergences in differential expression

To fully incorporate probability distributions, we propose a novel distance metric between subjects based on each gene *g* in the cytoKernel-sc approach. This metric is objective and can be easily tested for differential expression within a semi-parametric logistic regression framework. We first discuss the concept of divergence or distance between two probability distributions, followed by its implementation.

#### 2.3.1 Embedding of Conditionally Negative Definite (CND) via Hilbert Space

Consider a measure space (𝒳, 𝒜, *µ*), as conceptualized by Billingsley (2008); Callut et al. (2011), where 𝒳 is the sample space and 𝒜 the *σ*-algebra of measurable subsets, with *µ* representing a dominating measure. This framework delineates the set of all probability distributions 𝒫, where each distribution *P* maps elements of 𝒜 to the interval 𝒳. Central to this context is the Jensen-Shannon Divergence (JSD), a divergence measure *D*_*JS*_(·, ·) between two probability distributions *P*_1_, *P*_2_ ∈ 𝒫, defined as:

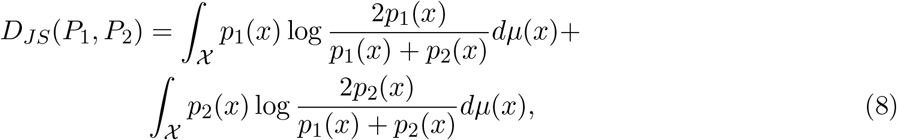

where *p*_1_ and *p*_2_ are the respective Radon-Nikodym derivatives of *P*_1_ and *P*_2_ with respect to *µ*.

In addition, the square root of the JS divergence, denoted here by 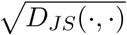, unlike divergences such as the Kullback-Leibler divergence (van Erven and Harremos, 2014), enjoys the properties of a true distance metric, *d*((·, ·) : identity, symmetry, and the triangle inequality (proof in Supplementary Materials). These properties enable JS divergence to effectively quantify the similarity between probability distributions, with smaller JS divergence values indicating higher similarity.

To maintain clarity, we use the same notations used in describing the data triplets {**z**_*i*_, **x**_*i*_, *y*_*i*_}, *i* = 1, …, *n* where *n* is the total number of subjects obtained from a case control study for feature *j* (omitting the gene index *g*). Without loss of generality, for each subject *i*, the expression of the observed data from the single cell vector **z**_*i*_ is conceptualized as a continuous random variable, symbolized by *Z*_*i*_ following the single cell data framework for JS divergence based distance metric construction (Seal et al., 2022). Now, *Z*_*i*_ is observed across *n*_*i*_ cells for subject *i*, represented as, 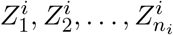.The cumulative distribution function and probability density function of *Z*_*i*_ are represented by *F*_*i*_ and *f*_*i*_, respectively.

For the sake of completeness, let 𝒫 be a convex set of probability measures defined on a measure space (𝒳, 𝒜, *µ*), where 𝒳 is the input sample space, and 𝒜 the *σ*-algebra of measurable subsets, with *µ* representing a dominating measure. Within this setup, the collection 𝒫 contains the probability distributions function *F*_*i*_ for *i* = 1, 2, …, *n*.

The dissimilarity between two subjects *i* and *i*^*′*^ in terms of the single cell distribution is quantified using a distance measure, denoted as 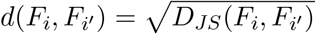, based on the following equation:

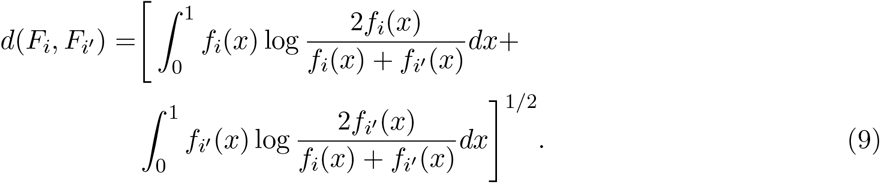

A high value of 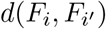 suggests a significant variance in the distribution or density between pairwise subjects *i* and *i*^*′*^, whereas a low value indicates similar distributions. Subsequently, a distance matrix between all the pairwise subjects, denoted as 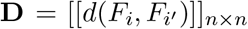, can be constructed. In practical scenarios, the density function *f*_*i*_ is not known a priori. Hence, we estimate it using Kernel Density Estimation (KDE), denoted as 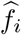, based on the observations 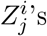 for *j* = 1, …, *n*_*i*_. The KDE is expressed as:

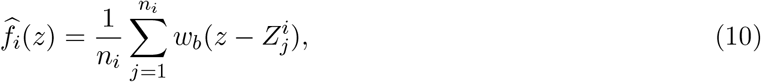

where *w*_*b*_ is a Gaussian kernel with a bandwidth parameter *h*, selected according to Silverman’s rule of thumb (Silverman, 1981). Utilizing these KDEs, 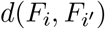 is approximated as:

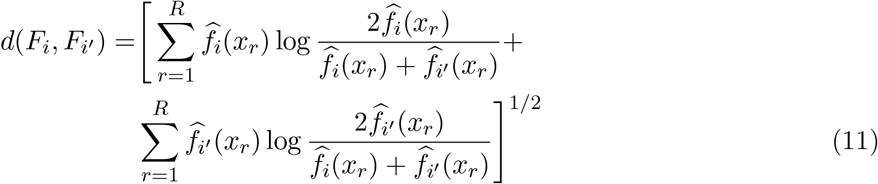

where *x*_*r*_, *r* = 1, …, *R* are grid points within the interval [0, 1]. In our simulations and empirical analyses, we observed that the estimates remain stable for sufficiently large values of *R*. We set *R* to 1024 and used evenly spaced grid points, ensuring that the estimated densities sum up to 1 through appropriate scaling.

In the supplementary materials, we define the Conditionally Negative Distance (CND) and show that, 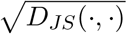 induces an embedding of the distributions in a real Hilbert Space. The Jensen Shannon Divergence can be described as the dot product of two probability distributions in a real Hilbert Space. That is, 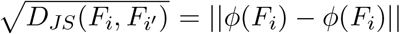, where *ϕ* is a function that maps the probability distributions in a Hilbert Space. A fundamental property used to prove the existence of the embedding is the notion of conditionally negative definite (CND) function. Further, (Topsoe, 2003) has shown that JS Divergence is Conditionally Negative Definite (CND).

### 2.4 Overview of Multivariate Distance Matrix Regression (MDMR)

For large number of subjects, semiparametric kernel regression offers computational advantages by calculating p-values based on the asymptotic distribution of the test statistic. However, for studies with small to moderate subject sizes, distance-based pseudo F tests, also referred to as Multivariate Distance Matrix Regression (MDMR), is preferred. MDMR relies on the resampling procedure to determine p-values. In both semiparametric kernel regression and MDMR approaches, the distance matrix must be converted to a kernel matrix.

We now briefly review Multivariate Distance Matrix Regression (MDMR) (McArdle and Anderson, 2001; Reiss et al., 2010; Zapala and Schork, 2012). Let 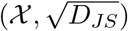 be a semi metric space and **Z** be a random object taking values in 𝒳. Suppose, we observe independent draws of single cell vector **z**_*i*_ for subject *i* = 1, 2, …, *n* and *D* = (*d*_*ij*_)_*n*×*n*_ denotes the sample dissimilarity (distance) matrix for subject pair (*i, j*), *i, j* = 1, 2, …, *n* such that, (*d*_*ij*_) can be intuitively written as,

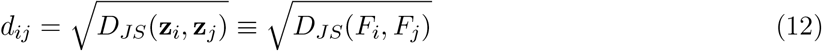

Define the double centered matrix following the notations used in the previous section, we write, *G* = *HAH* with *H* = *I* − *E/n* and *A* = −*D*^2^*/*2, where *I* denote an *n* × *n* identity matrix and *E* an *n* × *n* matrix where each element is equal to 1. The matrix *H* is a projection matrix, known as the centering matrix (Mardia et al., 1979)..

Following the notations used in the kernel-based regression, we write, **X** as an *n*×*q* covariate matrix. Let **X**^∗^ be the set of variables augmenting the binary outcome variable of interest **y** = (*y*_1_, *y*_2_, · · ·, *y*_*n*_)^*T*^ with all the covariates **X**, i.e., **X**^∗^ = [**y, X**] which can be expressed as a *n* × (*q* + 1) design matrix with corresponding projection matrix **H**_**X**∗_ = **X**^∗^(**X**^∗*T*^ **X**^∗^)^−1^**X**^∗*T*^. Here **H**_**X**∗_ is the traditional hat matrix.

MDMR assesses the association between **Z** and **X**^∗^ via the positive definite kernel matrix *G*. A pseudo F test statistic, as introduced by McArdle and Anderson (2001), is then represented by:

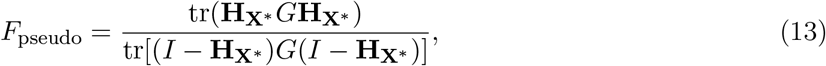

where the statistical significance is evaluated via permutation test (Tang et al., 2016; Zhang et al., 2022).

### 2.5 cytoKernel-psrF

In this section, we propose a new test statistic, *F*_sqrt_, a square root of the pseudo F statistic derived from McArdle and Anderson (2001); Li et al. (2009); Tang et al. (2016). In single cell studies, moderate to high correlations are observed among feature expression, i.e., gene expression for scRNAseq and protein marker expression for mass cytometry. To account for these correlated structure of the response variable (feature expression) and to boost power of the test, we define a new statistic following Shi et al. (2023).

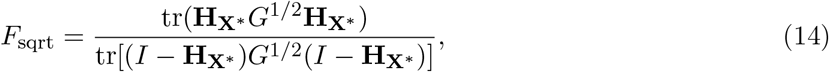

where the square root of a matrix *A* is denoted as *B*, satisfying *B* = *A*^1*/*2^. Let 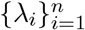 and 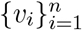 be the eigenvalues and eigenfunctions, respectively, of 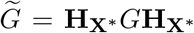. We define 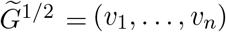 diag 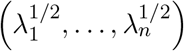

(*v*_1_, …, *v*_*n*_)^⊤^. The numerators of *T*_pseudo_ and *T*_sqrt_ can be intuitively represented as 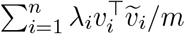 And 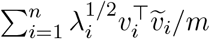, respectively, where **H**_**X**∗_ has eigenvalues equal to 1 and eigenfunctions 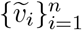. The square-root method potentially enhances the test efficacy by increasing the weight of significant factors when the response variables exhibit moderate correlation (Shi et al., 2023).

In the presence of covariates, we employ the conditional distribution of the residuals rather than augmenting the binary variable of interest and all the covariates.

For scenarios where confounding variables **X** exist, we adopt a strategy of resampling the residuals post-regression of **y** on **X**. The linear predictor in this regression model is represented as *γ***X**.

For cases where **y** is binary, the computation of the residual is performed via logistic regression, and the determination of the *p*-value is achieved through the application of the parametric bootstrap technique, as detailed by Davison and Hinkley (1997). The process involves the following steps:

- Perform a regression of **y** on **X**, leading to the maximum likelihood estimation (MLE) of *γ*, denoted as 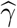. Calculate the residuals

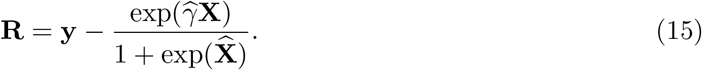

From these residuals, construct the observed square root of the pseudo-F statistic (*F*_*sqrt*_).
- For each resampling iteration, generate **y**^∗^ from a Bernoulli distribution with the success probability exp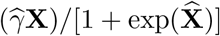. Conduct a regression of **y**^∗^ on **X** to obtain the MLE 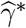 for *γ*. Calculate the permutation residuals

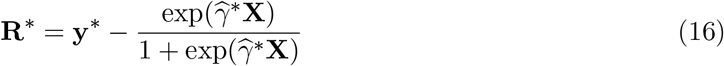

and form the resampled square root of the pseudo-F statistic *F*_*sqrt*_ based on **R**^∗^.
- Determine the final *p*-value by calculating the proportion of resampled *F*_*sqrt*_ statistics that exceed the observed statistic.

This resampling strategy effectively addresses the influence of confounder covariates and provides a robust approach to statistical analysis in the presence of binary variables and confounding factors.

## 3 Results

In our analysis of both simulated and real data, we focused on the cytoKernel-psrF variant of the cytoKernel method due to the small number of subjects in realistic sc-seq data settings. We will refer to cytoKernel-psrF as cytoKernel in the following simulations and real data sections. In the supplementary materials, we presented a comparative analysis among three distinct cytoKernel methodologies: cytoKernel-PB, cytoKernel-sc, and cytoKernel-psrF in the context of the SplatPOP benchmarking. By varying the number of subjects, we comprehensively assessed the performance of each method under different settings.

### 3.1 Simulations

#### 3.1.1 diffcyt

We analyzed the semi-simulated mass cytometry data from the designs implemented in Weber et al. (2019); Tiberi et al. (2022). These simulations, which incorporated spike-in signals into experimental data (Bodenmiller et al., 2012), were designed to evaluate the performance of the cytoKernel method. This approach preserved real biological data characteristics while embedding a known ground truth. Our evaluation focused on cytoKernel and two diffcyt methods based on limma and linear mixed models (LMM), both of which previously showed superior performance on these datasets.

We examined three datasets from Weber and Soneson (2019): the primary DS dataset and two variants with 50% and 75% diluted differential effects (Weber et al., 2019). Each dataset contained 24 protein markers, 88, 435 cells, and two groups across eight samples each. We implemented two cell grouping approaches: one using eight manually annotated cell types (Figure 1a) and another with 100 high-resolution clusters determined by unsupervised clustering method FLOWSOM (Van Gassen et al., 2015) (Figure 1b).

**Figure 1:**
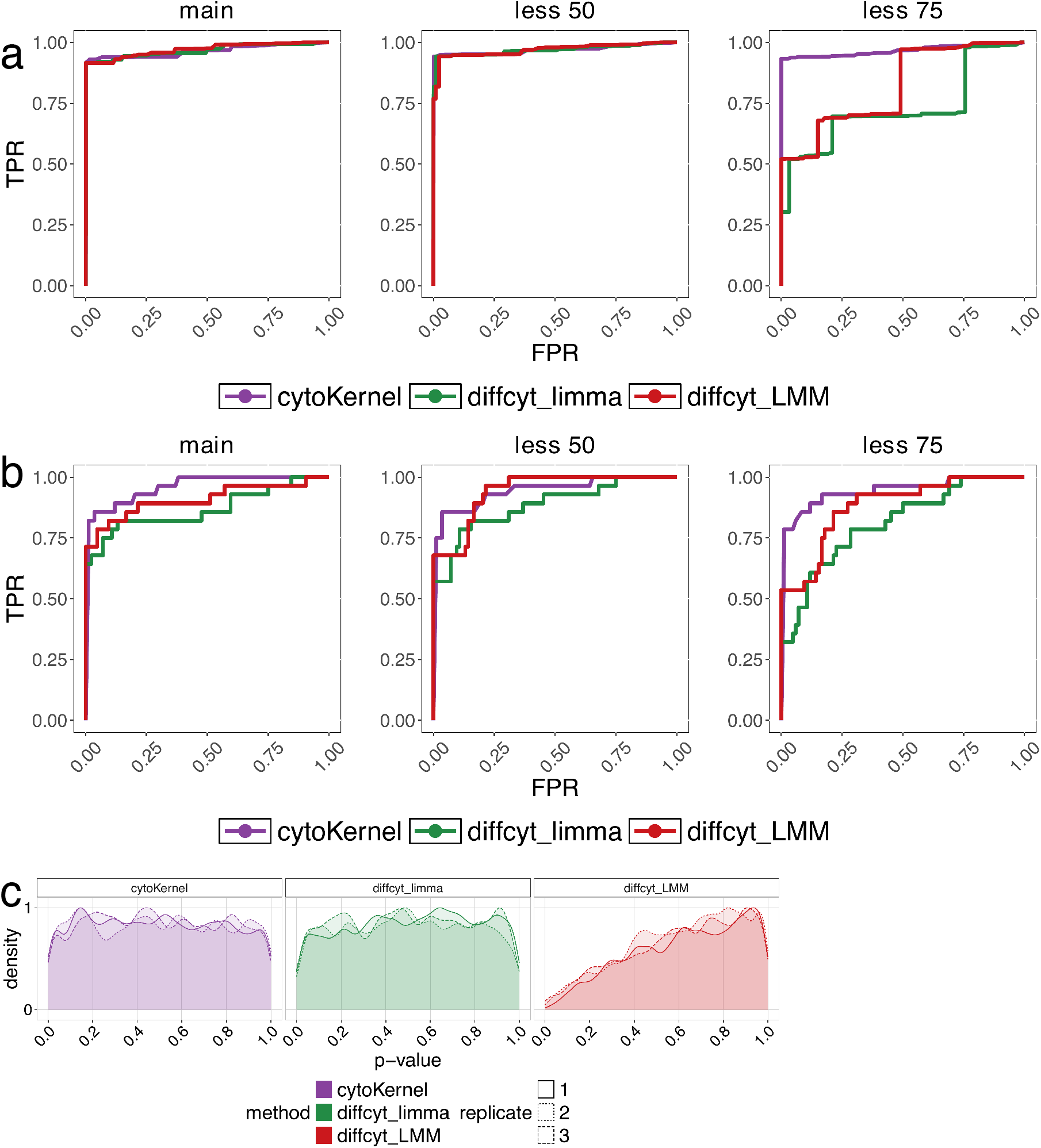
Receiver operating characteristic (ROC) curve in diffcyt non-null semi-simulated data. Each simulation consists of 88,435 cells and two groups of 8 samples each. ‘main’, ‘less 50’ and ‘less 75’ indicate the main simulation, and those where differential effects are diluted by 50% and 75%, respectively. We evaluated methods’ performance in terms of detecting DS for phosphorylated ribosomal protein S6 (pS6) in B cells, which is the strongest differential signal across the cell types in the dataset (Weber et al., 2019). Cells were clustered based on (a) manually annotated cell types and (b)unsupervised FLOWSOM clustering as in distinct simulation study (Crowell et al., 2020; Weber et al., 2019; Tiberi et al., 2022). (c) Density of raw *p*− values in diffcyt null semisimulated data (Weber et al., 2019; Tiberi et al., 2022). Each replicate represents a different null simulation. Each replicate consists of 88, 438 cells and two groups of eight samples each. Cells were clustered in an unsupervised manner (Weber et al., 2019).

The primary simulation study observed that the cytoKernel method exhibited a notably higher True Positive Rate (TPR) when cell-type labels were used. In contrast, when unsupervised clustering was applied, all methods showed comparable performance except 75% diluted differential effects where cytoKernel showed higher TPR (Figure 1a). Notably, as the magnitude of the differential effect decreased, the disparity in performance became more pronounced. Specifically, the diffcyt methods showed a significant reduction in TPR, whereas cytoKernel not only maintained a higher TPR (Figure 1b) but also effectively controlled the False Discovery Rate (FDR) (Supplementary Materials). This outcome indicates the robustness of cytoKernel in identifying even minor differential changes similar to distinct (Tiberi et al., 2022).

Furthermore, the study analyzed three replicated null datasets from Weber et al. (2019). These datasets consisted of 24 protein markers and 88, 438 cells distributed across eight cell types and were characterized by the absence of any differential effect. In these null scenarios, all evaluated methods, except for LMM, yielded *p*-values that were uniformly distributed, as shown in Figure 1c. This uniform distribution of p-values signifies the reliability and validity of the cytoKernel method in scenarios free of differential effects.

#### 3.1.2 muscat

We simulated droplet scRNA-seq data using muscat, as previously described in Crowell et al. (2020); Tiberi et al. (2022). The study involved five replicates simulating differential characteristics profiles, with 10% of genes in each cluster exhibiting differential characteristics. These characteristics span across various differential patterns as described in Korthauer et al. (2016); Tiberi et al. (2022):

- Differential Expression (DE) indicating a shift in the entire distribution.
- Differential Proportion (DP) indicating varied proportions in mixed distributions.
- Differential Modality (DM) contrasting a single-mode with a dual-mode distribution.
- Differential Modality and Means (DB) comparing a single-mode and dual-mode distribution with equal means.
- Differential Variability (DV), where two single-mode distributions with identical means but different variances were compared.

The simulations included 4, 000 genes across 3, 600 cells in three clusters, split into six groups with three subjects each, averaging 200 cells per subject per cluster. Our study examined how changing the number of cells per sample in each cluster influences the results. Specifically, we extended our simulations to include scenarios with 50, 100, 200, and 400 cells in each subject. This approach facilitated the evaluation of how variations in cell count influenced the sensitivity of the analysis. The results are compiled from five repeated simulations for each type of differential characteristic (DE, DP, DM, DB, and DV), with each type contributing an equal fraction. Further, we compared our simulation performance with IDEAS-PERMANOVA-S (Tang et al., 2016; Zhang et al., 2022).

The study examined six normalization techniques: Counts, Counts Per Million (CPMs), the logcounts method from scater (McCarthy et al., 2017), linnorm (Yip et al., 2017), BASiCS (Vallejos et al., 2015; Eling et al., 2018), and the residuals from variance stabilizing normalization (vstresiduals) (Hafemeister and Satija, 2019). These methods were evaluated alongside Poisson-based procedures from muscat, such as edgeR (Robinson et al., 2010) and limma-trend (Ritchie et al., 2015). These Poisson-based methods have been recognized for their effectiveness in differential analysis of scRNA-seq data (Crowell et al., 2020; Skinnider et al., 2021; Squair et al., 2021). In addition, scDD (Korthauer et al., 2016) based methods were assessed, as implemented in distinct (Tiberi et al., 2022). scDD utilizes a nonparametric method to detect distribution changes in scRNA-seq data, employing only the Kolmogorov–Smirnov test (scDD-KS). We omit the permutation approach (scDD-perm) due to its high computational cost.

CytoKernel maintained or surpassed the performance of leading methods such as edgeR.linnorm and limma-trend.logcounts in scenarios with 50, 100, and 200 cells as shown in the ROC (Receiver Operating Characteristic) curves (Figure 2). It demonstrated robust performance across all scenarios, effectively balancing statistical power with False Discovery Rate (FDR) control, as detailed in the Supplementary material. Overall, increasing cell counts generally improved the performance of all methodologies, particularly in detecting differential effects, as evidenced by higher true positive rates. Further analysis indicated that the performance of cytoKernel was minimally influenced by different normalization inputs, likely due to its inherent nonparametric design.

**Figure 2:**
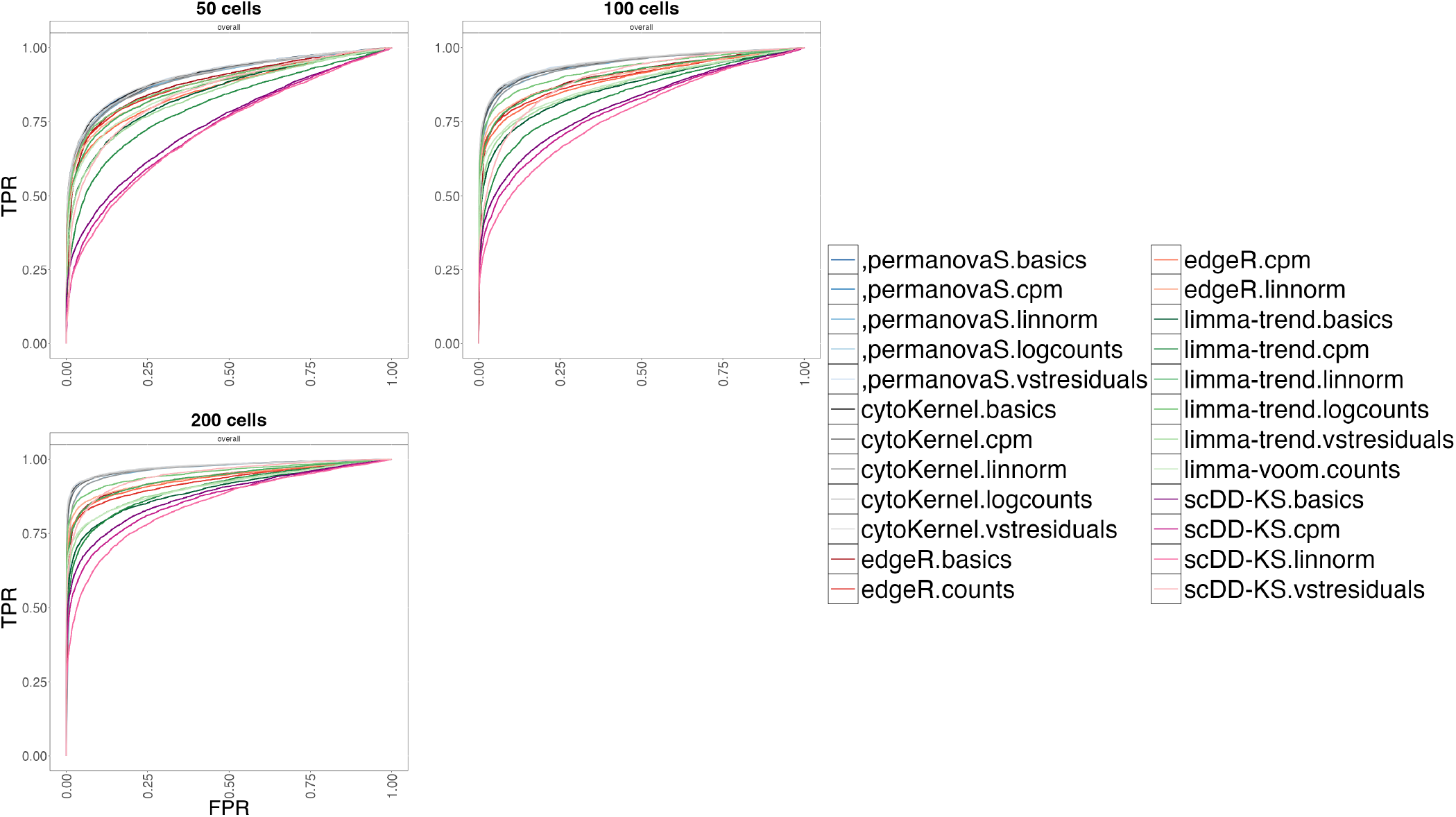
Receiver operating characteristic (ROC) curves generated using the distinct sensitivity (varying the number of cells) simulation design (Crowell et al., 2020; Tiberi et al., 2022). cytoKernel demonstrates better performance when varying the number of available cells. TPR versus FPR in muscat simulated data; with 50, 100 and 200 cells per cluster-sample combination, corresponding to a total of 900, 1800 and 3600 cells, respectively. Results are aggregated over the five replicate simulations of each differential type (DE, DP, DM, DB, and DV), contributing in equal fraction. Each individual simulation replicate consists of 4000 genes, three cell clusters and two groups of three samples each.

### 3.2 Real data

#### 3.2.1 Null experimental data

For evaluating false positive rates (FPRs) using real data, two scRNA-seq datasets under identical experimental conditions were analyzed, as previously examined in Tiberi et al. (2022). These datasets were not expected to exhibit differential characteristics. The focus of the analysis was on cytoKernel and Poisson-based (PB) methods, considering the high computational requirements and limited effectiveness of Mixed models (MM) methods (Crowell et al., 2020), as well as the elevated FDR noted in scDD models (Korthauer et al., 2016). The analysis included gene-cluster pairs with at least 20 non-zero cells in all subjects.

The first dataset, “Kang”, includes 10x droplet-based scRNA-seq data of peripheral blood mononuclear cells from eight Lupus patients, both before and after a 6-hour treatment with interferon-*β* (INF-*β*) (Kang et al., 2014). This dataset comprises 35,635 genes and 29,065 cells, manually categorized into eight cell types. Due to its outlier characteristics, one patient was excluded (Tiberi et al., 2022). The analysis concentrated on singlet cells and cells assigned to specific populations, considering only the control samples, which resulted in 11,854 cells and 10,891 genes. Three replicated datasets were then generated by randomly dividing the seven remaining control samples into two groups of three and four.

The second dataset, called “T cells”, is a Smart-seq2 scRNA-seq dataset encompassing 19,875 genes from 11,138 T cells obtained from the peripheral blood of 12 colorectal cancer patients (Zhang et al., 2019). Cells were sorted into 11 clusters utilizing igraph (Csardi and Nepusz, 2006). To create replicated datasets, the 12 patients were randomly split into two groups of six, forming three replicates.

In the analyses of the “Kang” dataset under null conditions, it was observed that the application of limma-trend, particularly when utilizing Counts Per Million (CPMs), resulted in elevated false positive rates (FPRs). The p-values derived from cytoKernel indicated a slight increment towards zero. In contrast, edgeR and limma-voom demonstrated more conservative properties, thereby providing enhanced control over FPRs, as depicted in Figure 3. The “T cells” dataset is presented in the Supplementary Materials. With regards to normalization methods, both linnorm and BASiCS were found to generate the most conservative p-values, consequently leading to the lowest false positive rates observed in the study.

**Figure 3:**
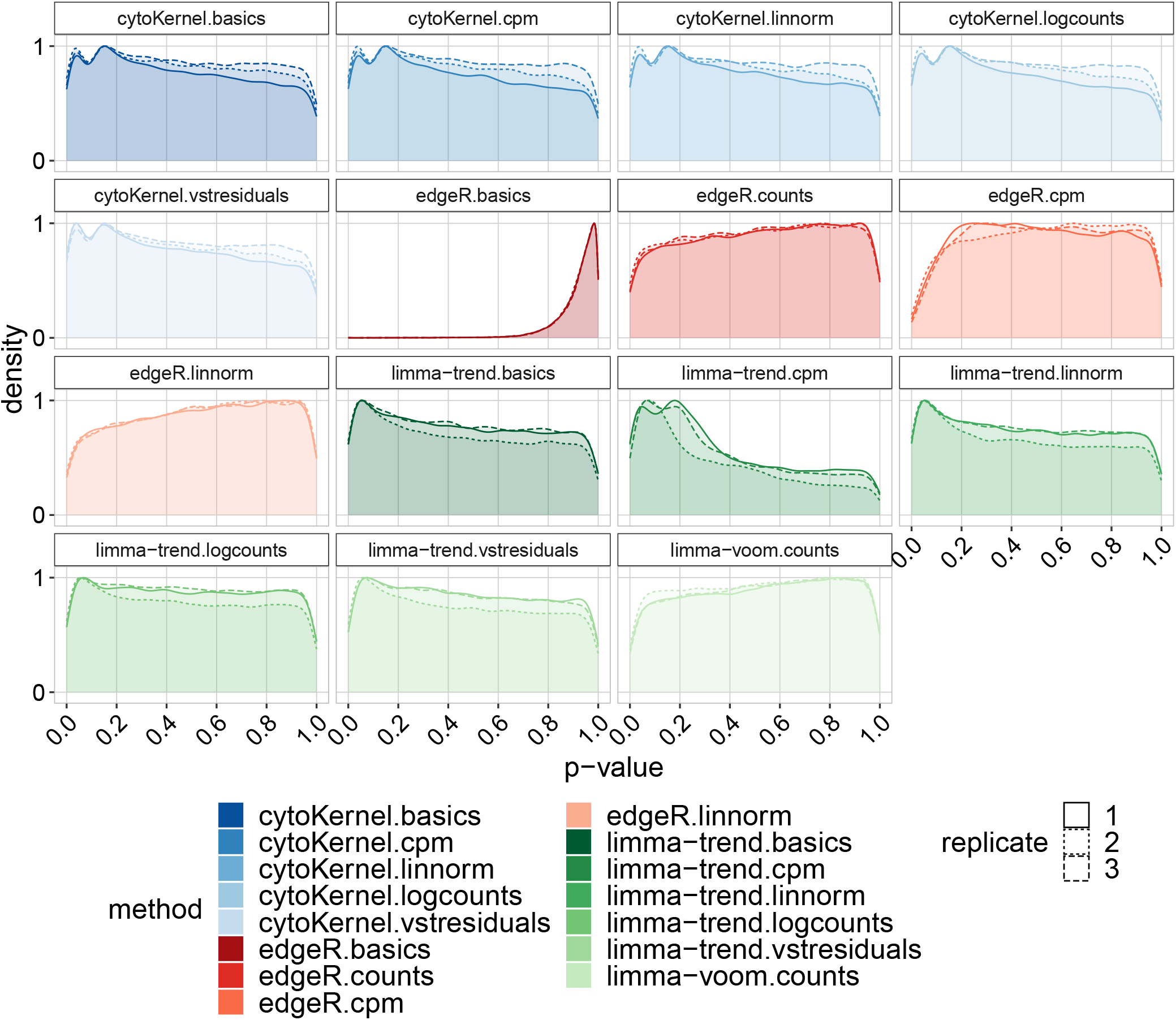
Density of raw p-values in null Kang Lupus data, comprising 11,854 cells across eight clusters.. Each replicate in these datasets represents a random division of samples into two groups, highlighting the distribution of p-values obtained. The ‘cytoKernel’ method demonstrates an almost-uniform distribution of null p-values.

#### 3.2.2 Bacher COVID-19 data

In a study performed by Bacher et al. (2020), a detailed statistical analysis was conducted to identify differentially expressed genes in central memory and Type I interferon-gamma (IFNG) T-cell subpopulations across patients with varying severity of COVID19. In the study, the Benjamin-Yekatueli procedure (Benjamini and Yekutieli, 2005) was utilized for False Discovery Rate (FDR) control, specifically tailored for dependent gene sets. This statistical approach enhances the accuracy of identifying differentially expressed genes by accounting for dependencies within the data. Upon examining the gene-cluster combinations flagged by cytoKernel using the criteria: adjusted *p* − *value <* 0.1, we identified several significant genes. UMAP visualizations effectively delineated these subpopulations in correlation with disease severity, (mild-moderate and severe). The primary focus of the study was to identify differentially expressed genes from subpopulations of interest and to visually inspect the empirical distributions of crucial genes like IL2 and IFNG (Supplementary material). This approach aimed to uncover non-canonical differential expression characteristics associated with COVID-19 severity. Additionally, in Figure 4, concordance plot using the upsetR package provided insight into gene expression overlaps and divergences across different T cell clusters, enhancing the understanding of immune responses to SARS-CoV-2. It details how differential expression patterns are compared across identified cell subpopulations, particularly between mild-moderate and severe COVID-19 cases, using a significance threshold of *p <* 0.1. The analysis reveals that the central memory and Tfh-like subpopulations show a higher incidence of differential expression patterns compared to Type 1-IFN-signature and Cytotoxic-Th1. Interestingly, the Cycling subpopulation is characterized as the most conservative in terms of differential pattern identification. Additionally, a significant consistency in differential expression is noted across memory subpopulations, specifically Transitional and Central memory.

**Figure 4:**
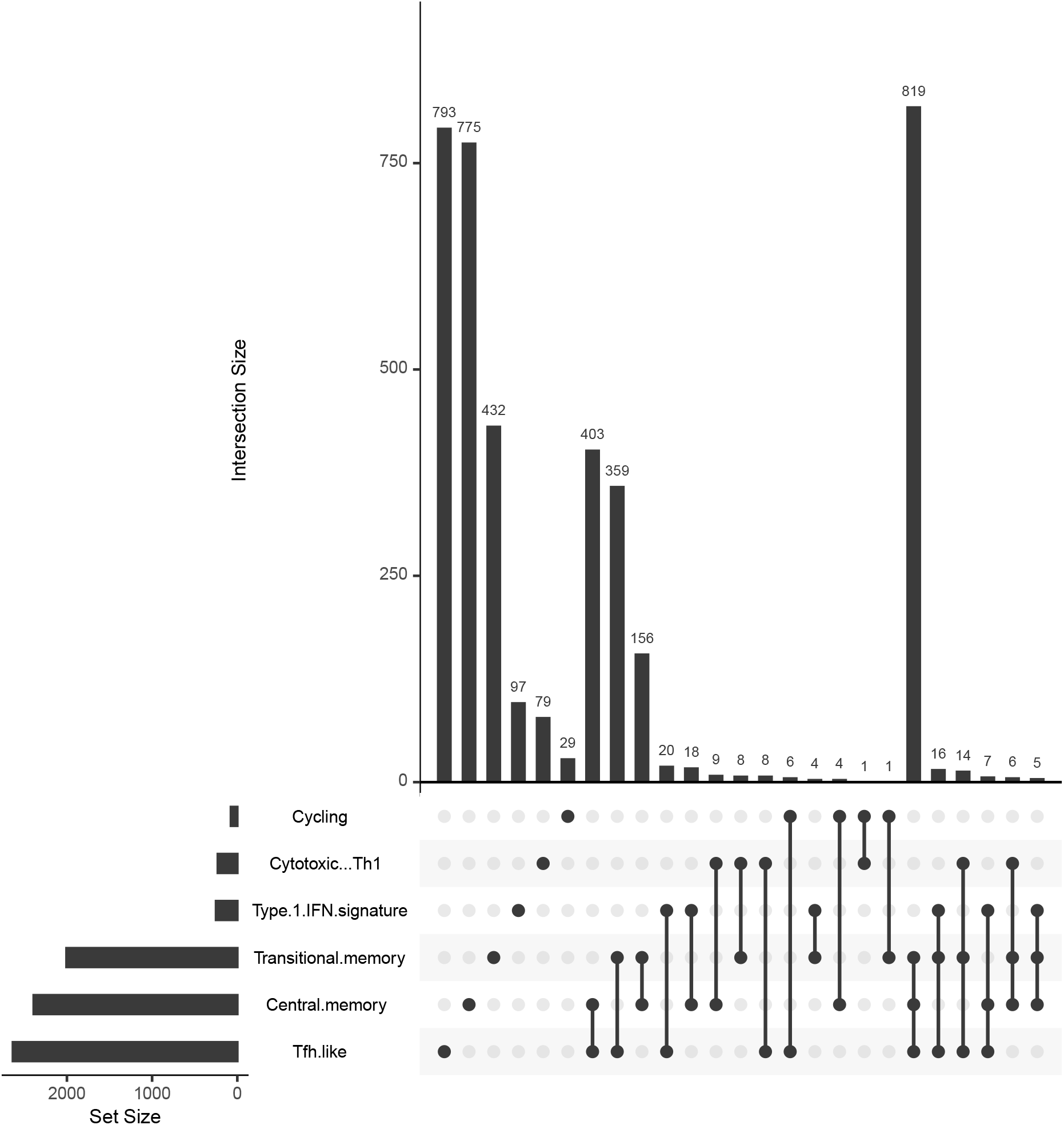
Analytical Visualization of Differential Expression Intersections across identified cell subpopulations. The graphical representation, generated via the UpSetR R package, compares the intersections of differential expression patterns identified by cytoKernel in the Bacher COVID-19 dataset. This comparison, focusing on mild-moderate versus severe subjects under an adjusted p-value criterion of < 0.1, highlights that the central memory and Tfh.like identify a greater number of differential expression patterns compared to Type 1-IFN-signature and Cytotoxic-Th1 subpopulations. Notably, the Cycling subpopulation emerges as the most conservative in identifying differntial patterns. Furthermore, the memory subpopulations (Transitional memory and Central memory) display a notable consistency across Memory subpopulation sets.

## 4 Discussion

Traditionally, single-cell sequencing (sc-seq) datasets have faced limitations due to high sequencing costs and technical constraints. This has resulted in data being collected from various cell types but from only a few subjects. Consequently, the focus has been on identifying differential expression across cell types. There has been less emphasis on comparing multiple subjects between case and control groups. However, with the decreasing costs and improved accessibility of sc-seq datasets, there is a growing availability of vast case-control study datasets, particularly in complex human diseases research. To address the needs of these emerging datasets, we have developed a novel method, ‘cytoKernel’, tailored for differential expression analysis in single-cell data. ‘cytoKernel’ utilizes a robust multi-subject full-distribution kernel embeddings framework, designed to identify differential patterns between groups of distributions, especially effective in scenarios where mean changes are not evident.

Our method has demonstrated superior performance in extensive benchmarks against both simulated and experimental datasets from scRNA-seq and mass cytometry, offering better control over false-negative and false discovery rates and identifying more differential expression patterns than traditional methods. Notably, it shows higher statistical power than pseudo-bulk methods and other statistical frameworks, like Bayesian hierarchical framework based scDD. ‘cytoKernel’ also adeptly adjusts for sample-level, cell-cluster-specific covariates, including batch effects. The versatility of ‘cytoKernel’ extends beyond identifying differential patterns, and its performance remains consistent across various normalization approaches. This method, therefore, marks a significant advancement in the analysis of high-throughput single-cell data, particularly in biomedical research involving complex disease groups.

Data-driven methods, particularly those identifying differential expression, are vital in guiding experimental research focused on complex diseases such as Lupus and COVID-19. These approaches are essential in genetic research, where discerning a precise list of key genes is far more beneficial than compiling an extensive yet less relevant gene catalog. The cytoKernel method exemplifies this utility with its nonparametric flexibility, strong adherence to controlling type-I error rates, and ability to detect crucial differential patterns of the genes. This makes it highly suitable for differential expression analyses in case-control studies with multiple subjects in each group. By efficiently isolating significant genes in these areas, cytoKernel provides a robust foundation for deeper experimental inquiry, enhancing our understanding of these complex disease conditions.

## 5 Acknowledgments

TG was supported by NSF SES 2149492 and the Grohne-Stepp Endowed Fund from the University of Colorado Cancer Center. RMB, VGL, and EWYH were supported by the Jeffrey Modell Foundation Translational Research Program and NIH NIAID R01 5R01AI174303.

## Notes

### Competing Interest Statement

The authors have declared no competing interest.

